# Diapause and Developmental Arrest as Drivers of Population Resilience Through Stress

**DOI:** 10.64898/2026.07.19.739441

**Authors:** Galayna Baur, Elizabeth Bone, Heather Moore, Logan Elliott, Alexis Ham, Molly Ripper, Andrea Scharf

## Abstract

Dauer formation and L1 arrest are stress-responsive developmental strategies that enable *Caenorhabditis elegans* to survive unfavorable conditions. These responses are regulated by environmental cues, including food availability and pheromone signals that communicate population density. However, how dauer entry, L1 arrest, and density-dependent signaling collectively influence long-term population dynamics remains poorly understood. In this study, populations of *daf-22* and *daf-16* mutants with impaired dauer formation, starvation arrest, and pheromone signaling were compared under control and starvation stress. Measurements of developmental stages were used to evaluate how genotype influenced population growth, developmental stage composition, starvation response, and recovery over time. This population-level approach links individual developmental decisions and inter-organismal communication to broader patterns of persistence and population change. The results show that *daf-16* and *daf-22* mutant populations differed from wild type in their recovery ability following nutrient deprivation as well as in the stage distributions within the population. These findings suggest that dauer signaling contributes broadly to population persistence by coordinating developmental arrest, reproduction, survival, and recovery. Overall, this work supports the interpretation of dauer formation as a larger population level survival program rather than a single isolated developmental outcome.

## INTRODUCTION

Biological systems consist of various levels of biological organization, and each level has emergent properties. Emergent properties are system-level features that arise through interactions among lower-level components and cannot be attributed to any single component in isolation (Reuter et al., 2005; Ponge, 2005). Biological organization is hierarchical, with each level giving rise to properties that are not present in its individual components. Interactions among molecules produce cellular functions, interactions among cells generate the structure and function of tissues and organs, and coordinated organ systems support the physiology and behavior of whole organisms.

Organisms then interact with one another and their environment to produce population-level properties, such as growth, density dependence, and persistence. At still higher levels, interactions among populations and their physical environment generate ecosystem-level processes, including community structure, energy flow, and nutrient cycling. Thus, emergent properties arise progressively through interactions among components at each level of biological organization (Begon et al., 2006). It is here in the levels of populations and ecosystems that we see the emergent property of population dynamics (Pelletier et al., 2011). Population dynamics are the changes in population density and developmental stage distribution of a defined population over a time window (Turchin, 2003). By investigating population dynamics, we are able to scale the traits of an individual such as life history traits, interactions, and movement to patterns of their resident populations, such as variance, size, and distribution (Benton et al., 2006; Grimm & Railsback, 2005; DeAngelis & Grimm, 2014). Because organisms exist within the context of both their population and environment, understanding biological responses requires consideration of population-level patterns alongside the traits and behaviors of individuals (Benton et al., 2006; Grimm & Railsback, 2005; DeAngelis & Grimm, 2014). This raises the question, how does an individual shape the population it lives in? How does the population shape the individual?

Understanding how genetic and environmental factors interact to shape population dynamics is a central question in biology. Individuals and populations persist within fluctuating environments that may be further disrupted by acute or sustained stress. How individual traits and genetic variation influence population dynamics and collective responses to these stressors remains poorly understood (Scharf et al. 2022).

The nematode *Caenorhabditis elegans* provides a powerful model for studying the relationship between genetic regulation and population behavior for two key reasons. 1) Under favorable conditions, *C. elegans* progresses through four larval stages before reaching adulthood (Figure 1) (Altun & Hall, 2009). Environmental conditions can redirect this developmental trajectory at distinct points in the life cycle (Hu et al., 2006). Larvae that hatch in the absence of food arrest in the L1 stage and resume development when food becomes available (Figure 1) (Baugh, 2013). Under later adverse conditions, including food limitation, elevated population density, and high temperature, larvae may instead enter the dauer stage (Figure 1) (Hu et al., 2006, Fielenbach & Antebi, 2008). Dauer formation is a collective-driven decision regulated by conserved signaling pathways that integrate environmental cues with pheromone-mediated communication among individuals (Hu et al., 2006, Fielenbach & Antebi, 2008). Together, L1 arrest and dauer represent distinct forms of developmental plasticity that promote survival during environmental stress (Baugh 2013, Hu et al., 2006, Fielenbach & Antebi, 2008). 2) Scharf *et al*. 2022 established a semi-continuous laboratory population system in which *C. elegans* and its bacterial food source were maintained through repeated cycles of controlled culling and feeding (Scharf et al., 2022). By tracking population abundance and developmental stage composition over time, this platform enables individual life-history traits to be linked to emergent patterns of population growth, structure and persistence.

**Figure 1.**
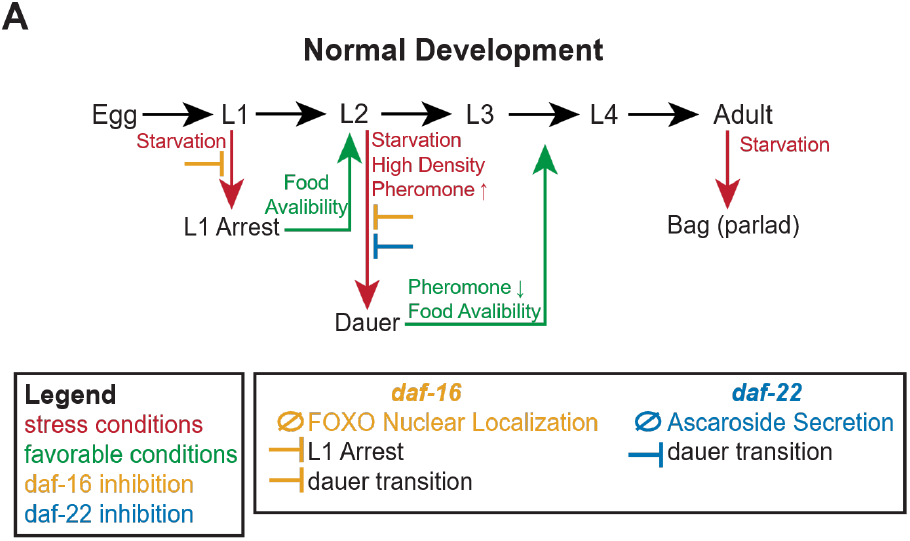
Experimental framework linking developmental plasticity to population dynamics. (A) Development of *C. elegans* with stress-resistant stages including blockage of the developmental transitions induced by *daf-16* and *daf-22* mutations. Arrows indicate developmental transitions, blunt headed arrows indicate inhibition of developmental transitions by genetic mutation.

The developmental responses of individual worms are shaped by genes that coordinate environmental communication with intracellular stress signaling. By linking environmental perception to developmental plasticity, these genes provide a potential mechanism through which individual-level traits scale to influence population dynamics. Among these genes, *daf-22* is required for the peroxisomal β-oxidation steps that generate density-dependent ascaroside pheromones involved in dauer signaling (Golden & Riddle, 1985; Butcher et al., 2009), whereas *daf-16* encodes a FOXO transcription factor that integrates insulin-like signaling to regulate development, dauer formation, metabolism, stress resistance, starvation survival, and longevity (Gottlieb & Ruvkun, 1994; Ogg et al., 1997; Baugh & Sternberg, 2006; Murphy & Hu, 2013) (Figure 1).

This allows us to dissect the connections between stress resistance, developmental decisions, inter-individual communication, and population persistence (Vowels and Thomas 1992; Ailion and Thomas 2000; Park et al. 2017; Gottlieb and Ruvkun 1994; Larsen et al. 1995; Tissenbaum and Ruvkun 1998; Isik et al. 2016). The molecular mechanism underlying dauer formation and stress responses are well characterized (Fielenbach & Antebi, 2008), however, it remains unclear how these pathways influence populations especially over many generations. Particularly, how mutations in developmental arrest and diapause abilities alter the timing, magnitude, and structure of population growth, persistence under fluctuating environmental conditions, and resource deprivation have not been systemically studied. To fill this gap, we investigate how mutations in key dauer-related genes affect population dynamics in *C. elegans* under non-stressed and food deprivation conditions. By combining population measurements across a series of timepoints with stage specific analyses and statistical modeling, we were able to quantify how genotypes influence population size, development, and response to environmental changes.

Here, we used a long-term laboratory population system (Scharf et al., 2022) to determine how genetic differences in developmental plasticity and density-dependent communication influence population dynamics under baseline and food-deprivation conditions. We compared wild-type N2, *daf-16* and *daf-22* populations by measuring abundance, developmental stage composition, starvation response and recovery over time. Genotype-dependent differences emerged across several features of population growth, structure and recovery, indicating that alterations to individual developmental and signaling pathways can influence population-level responses to environmental stress.

## METHODS

### STRAINS AND MAINTENANCE

*Caenorhabditis elegans* strains used in this study were N2 (wild type, WT), *daf-22* (ok693), *daf-16* (mu86). All strains were obtained from the Caenorhabditis Genetics Center (CGC) which is funded by NIH Office of Research Infrastructure Programs (P40 OD010440). Worms were maintained at 20°C on nematode growth medium (NGM) plates seeded with OP50 *Escherichia coli* following standard protocols (Brenner, 1974; Stiernagel, 1999).

### EXPERIMENTAL DESIGN/ TREATMENTS

#### Synchronization

All experiments were started with age synchronized *C. elegans*. First, worms were washed from mixed age NGM culture plates with M9 buffer and pelleted. Next, the worm pellet was resolved in bleach solution [2 mL 4 M NaOH, 4 mL household bleach, and 4 mL dH₂O] and pelleted at 3000 x g in a centrifuge, this was repeated to a total of two bleachings. Bleaching terminated all stages of worms other than the eggs. Then, worms were washed three times in sterile ddH2O, resolved in 1 mL of sterile M9 [3 g KH₂PO₄, 6 g Na₂HPO₄, 5 g NaCl and 1 mL 1 M MgSO₄, 1 L dH₂O], and incubated on a shaker at 100 rpm and 20°C overnight (Stiernagel, 1999).

#### Population Dynamics

Populations were maintained at 20°C and on a 100-rpm shaker; all culling/ feeding was done under a sterile hood with aseptic technique according to Scharf et al., 2022. All assays were completed in parallel with 3-5 biological replicates per condition, per experiment. Initiating Populations (Scharf et al., 2022): On day one, synchronized worms were introduced into a culture flask containing 10 mg OP50 *E. coli*, and 5 mL sterile S-medium [100 mL S-basal [5.84 g NaCl, 6.8 g KH₂PO₄, 1 mL of Cholesterol in Ethanol (5 mg/mL), 1 L dH₂O] and left overnight to grow. 5 µL samples were then counted a minimum of five times per population and averaged to obtain the worms/µL concentration. Population flasks were initiated with 100 larvae in 5 mL sterile S-medium, and 10 mg OP50 *E. coli* then left to grow for 48 hours. Next populations were fed with 0.5 mL of OP50 in S-Medium (20 mg/mL) and grown for 24 hours. Culturing: Once every 24 hours for the culture period, 10% (0.5 mL) of the populations were culled followed by refeeding with an equivalent amount (0.5 mL) of OP50 in S-Medium (20 mg/mL). Culled worms were analyzed by counting multiple samples of 5-10µL, acquiring the average number of eggs, larvae, and adults per population.

#### Deprivation Dynamics

Deprivation dynamics were tested in parallel with continuously fed population dynamics experiments as controls. Populations were initiated and maintained following the same protocol as for population dynamics, with the addition of a deprivation period. Starting on day 20, the deprivation period consisted of no culls or feeds and lasted till refeeding on day 40 for the short experimental groups (Figure 5A) and day 50 for long the long experimental groups (Figure 6A). Each population was culled and fed every 24 hours for 20 days following the deprivation according to the population dynamics procedure (Scharf et al., 2022).

#### Liquid Culture L1 Arrest Assay

Synchronized worms were placed into culture flasks with S-medium at a concentration of 1000 worms/mL. Recovery was scored from this arrest flask every 5 days by transferring 10 µL worms onto seeded NGM plates as well as into 250 µL S-medium with 20mg/mL OP50 as food in a well plate. The worm’s recovery was scored after 72 hours (Hibshman et al., 2021).

#### SDS Assay

Samples from population dynamics culls were placed on unseeded NGM plates and treated with 1% SDS for 10 minutes with gentle agitation to terminate all worms without the thickened dauer cuticle before live worms were visually scored and counted as dauers (Karp, 2018). The dauer assay used showed no false positive dauer counts and had an approximately 6% false negative report when used on daf-2 dauer constitutive worms grown at 25°C (n=68 total larvae with 64 surviving larvae).

## ANALYSIS

Statistical analyses were performed in R (v4.5.2) using rstatix, lme4, and emmans packages. Plots were generated in R. Final figures were assembled and formatted, and schematics were created, using Adobe Illustrator (version 30.6; Adobe Inc.). Population trajectories were characterized using summary metrics representing the end of initial exponential growth (first peak), the highest and lowest post-peak population sizes (maximum and minimum respectively), the mean population size across the experimental period, and the time to each metric. Genotypic differences were evaluated using non-parametric tests owing to deviations from normality. Overall differences among genotypes within each condition were assessed using Kruskal-Wallis test (Kruskal and Wallis, 1952). When significant differences were detected, pairwise comparisons were performed using Wilcoxon rank-sum test (Wilcoxon, 1945) with Benjamini-Hochberg (BH) correction to control false discovery rates (Benjamini and Hochberg, 1995). Standard Error of the Mean (S.E.M.) was reported on all summary statistics and visualized as shaded areas in population dynamic graphs. Pairwise statistical results were reported using adjusted p-values. To account for repeated measurements and hierarchical structure within the data, linear mixed-effects models were employed. Models were fitted using restricted maximum likelihood with genotype and condition as fixed effects and replicate identity included as a random intercept term. Both condition-specific models and a full interaction model (genotype x condition) were evaluated. Post hoc pairwise comparisons of estimated marginal means were conducted using emmans package with BH-adjusted p-values. Empirical distribution differences between genotypes were further assessed using probability-probability (PP) plots. For each comparison, empirical cumulative distribution functions were computed and plotted for WT and mutant populations. Deviations from the identity line were interpreted as differences in distributional behavior between genotypes. All statistical tests were two-sided, and a significance threshold of p < 0.05 was used unless otherwise stated.

## RESULTS

### GENOTYPE DIFFERENCES IN POPULATIONS UNDER FLUCTUATING CONDITIONS

We first characterized population dynamics under baseline (control) conditions, where populations were fed and culled every 24 hours. WT populations exhibited a first peak on approximately day 18, before the population size stabilized; fluctuating around the average number of worms and remaining consistent for long term population persistence (Figure 2A, Table 1, Table 2). In contrast, once *daf-22* populations reached a delayed first peak, they appeared to steadily decline, potentially leading to extinction for longer term populations (Figure 2B). *daf-16* shows similar growth to WT populations in the initiation phase, but most resemble *daf-22* in the maintenance phase with a steadily declining population (Figure 2C). To assess how similar the distribution of population dynamics of the three genotypes were, we used PP-plots. These plots show that *daf-22* fall outside a 95% CI, shaded region, when compared to WT whereas *daf-16* fall predominantly inside the shaded 95% CI (Figure 2D, E). Extinction was not observed in any WT populations whereas 9% of *daf-22* and 10% of *daf-16* populations failed to persist under control conditions and went randomly extinct (Figure 2F). To determine whether broadly similar population trajectories differed in specific features, we next compared summary statistics describing population growth and decline. The first population peak of WT was significantly smaller than both dauer mutants, *daf-22* (p < 0.001) and *daf-16* (p < 0.01) (Figure 2G, Table 1). The maximum population size, the highest number of worms after the first peak, there was no significant difference in the number of worms, however we observed that *daf-16* populations reaching their max significantly before both WT (p < 0.05) and *daf-22* (p < 0.01) populations (Figure 2H, K, Table 1). There was no significant difference in the minimum population size, with all genotypes reaching their minimum at approximately the same time (Figure 2I, L, Table 1, Table 2). WT and *daf-16* reached their respective population peaks at approximately the same time on day 18, with *daf-22* following significantly behind on day 32 (p < 0.0001) (Figure 2J, Table 2). While there were differences in the average number of worms during the first twenty days of the experiment (Figure 3A), there were no significant differences in the average population size between all genotypes after the initiation phase (Figure 3B, C). Mixed-effects analysis further revealed no significant pairwise differences following multiple comparison corrections. This indicates that diapause-ability genotypes do not influence population size under non-stressed conditions. These results indicate that under non-stress conditions there are differences in population persistence of diapause-related mutants including a higher extinction risk.

**Figure 2.**
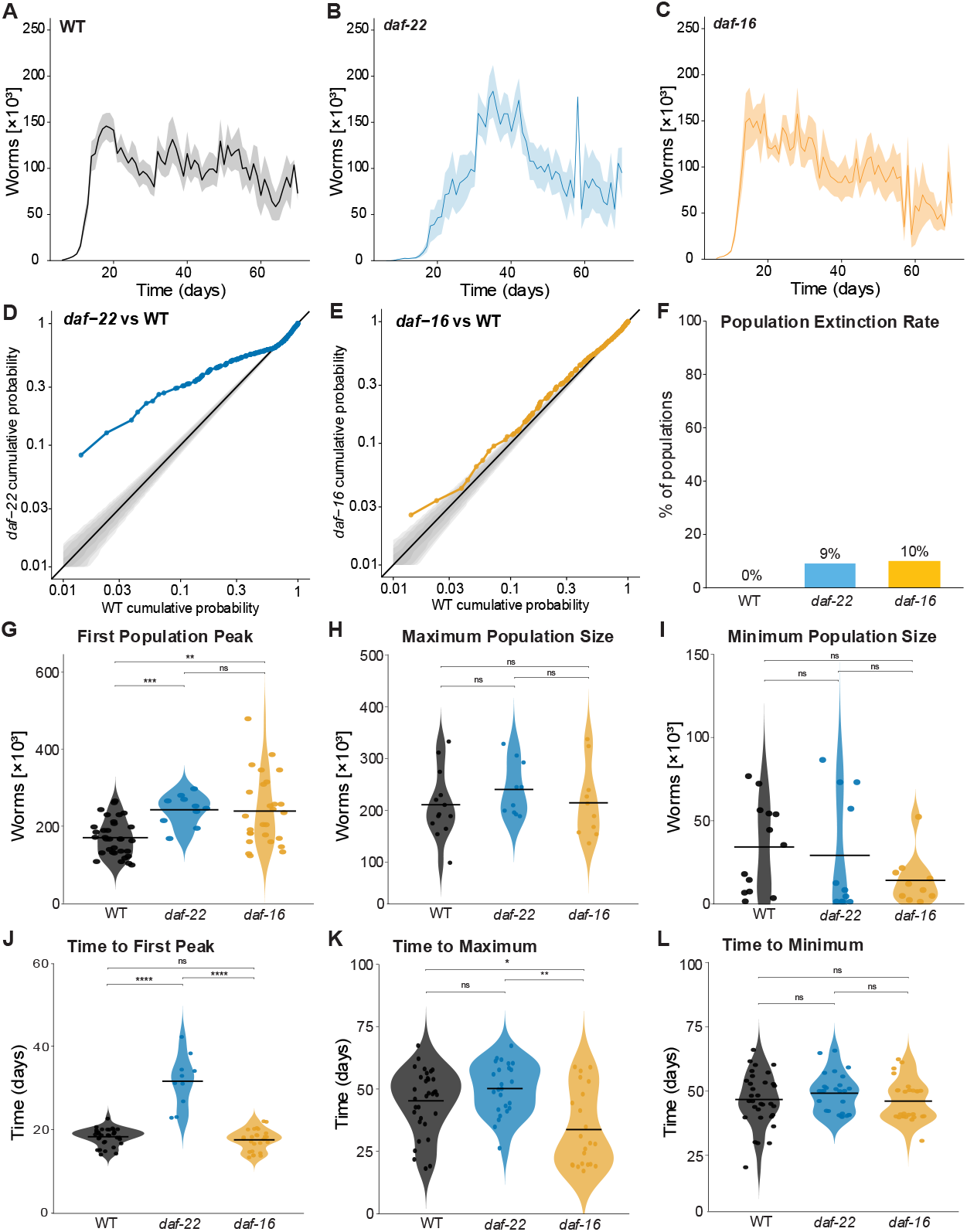
Genotypic differences in population dynamics emerge under control conditions. (A, B, C) Population dynamics graph of mean total population size over time. Values represent mean ± S.E.M. across biological replicates, with technical replicates averaged prior to analysis, WT is shown in black (n=15 populations), *daf-22* is shown in blue (n=13 populations). *daf-16* is shown in yellow (n=12 populations). Population counts are expressed as worms x10^3^. (D, E) PP plots comparing empirical cumulative distribution functions of WT and mutant populations. Axes are in log scale, the black line indicates identical distributions, and the shaded region represents a 95% confidence interval; deviations reflect differences in population dynamics. (F) Rate of extinction across genotypes. Extinction was defined as 15 or more days with no detected worms in population counts; the rate was defined as proportion of populations reaching extinction within each condition. (G-I) Population dynamics summary statistics (first peak, max, min) across genotypes: each point in violin plots represents a biological replicate, grey lines represent means. Wilcoxon rank pairwise test for significance. (J-L) Time taken by populations to reach summary statistics across genotypes. Each point represents a biological replicate, grey lines represent mean, Wilcoxon rank pairwise test for significance. Significance: p< 0.05= *, p< 0.01=**, p< 0.001=***, p< 0.0001= ****; not significant = ns.

**Figure 3.**
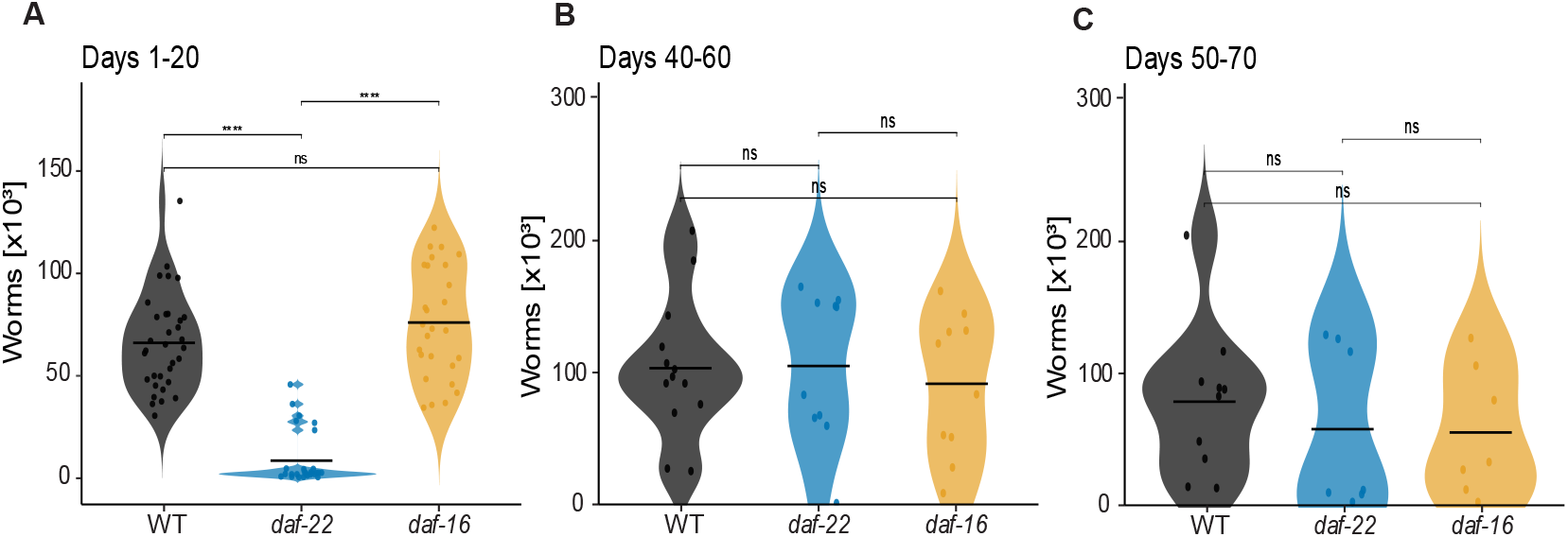
Genotype altered mean population size only during the initiation phase across 20-day intervals. Violin plots show the distribution of mean population sizes for each genotype during the indicated 20-day intervals. Each point represents one independent biological replicate population, and grey horizontal lines indicate group means. Sample sizes were n= 15 populations for WT, n=13 populations for *daf-22*, n=12 populations for *daf-16* at each interval. Pairwise differences among genotypes within each interval were assessed using two-sided Wilcoxon rank-sum tests, with P values adjusted using the Benjamini–Hochberg procedure. Significance: p< 0.05= *, p< 0.01=**, p< 0.001=***, p< 0.0001= ****; not significant = ns.

**Table 1.** Population Summary Statistics in a Fluctuating Environment.

| genotype | average | ± SEM | peak | ± SEM | max | ± SEM | min | ± SEM |
| --- | --- | --- | --- | --- | --- | --- | --- | --- |
| WT | 106008 | 11576 | 186231 | 14973 | 210969 | 17905 | 33842 | 7527 |
| *daf-22* | 93766 | 12306 | 242545 | 12649 | 240380 | 16448 | 27900 | 10633 |
| *daf-16* | 92662 | 14046 | 247400 | 37383 | 214620 | 22035 | 12875 | 4793 |

**Table 2.** Population Summary Statistics Timing in a Fluctuating Environment.

| genotype | time to peak | ± SEM | time to max | ± SEM | time to min | ± SEM |
| --- | --- | --- | --- | --- | --- | --- |
| WT | 18.30 | 0.36 | 45.24 | 2.20 | 46.73 | 1.79 |
| *daf-22* | 31.60 | 1.92 | 50.16 | 2.04 | 49.19 | 1.46 |
| *daf-16* | 17.56 | 0.47 | 33.73 | 3.26 | 46.11 | 1.51 |

## STAGE DISTRIBUTIONS REVEAL DEVELOPMENTAL DIFFERENCES IN MUTANT STRAINS

WT served as our control genotype with normal diapause abilities. *daf-16* is both dauer defective and defective in somatic L1 arrest because loss of functional DAF*-*16/FOXO prevents the transcriptional response required to execute dauer formation and arrest postembryonic development during starvation (Gottlieb & Ruvkun, 1994; Baugh & Sternberg, 2006). Under dauer-inducing or starvation conditions, functional DAF-16 normally localizes to the nucleus, where it regulates developmental-arrest and stress-response genes (Lee et al., 2001; Baugh & Sternberg, 2006). By contrast, *daf-22* is defective in the production of short-chain, density-dependent ascaroside pheromones and is therefore unable to signal dauer entry efficiently within a population, but it retains the ability to respond to exogenous dauer pheromone and is not known to be defective in L1 arrest (Golden & Riddle, 1985; Butcher et al., 2009; Artyukhin et al., 2013). Dauer abilities were confirmed with stage distribution counts showing that WT populations had dauers (Figure 4A) whereas there were no dauers in *daf-22* or *daf-16* populations whereas (Figure 4B, C). Next, we compared the survival rate of animals after extended L1 arrest between WT, *daf-22* and *daf-16* animals in liquid media. The rate of L1 arrest recovery was significantly decreased in *daf-16* individuals in comparison to WT animals at all measurement points (p < 0.01) (Figure 4D). With non-significant differences between WT and *daf-22* worms at all points (p > 0.05) aside from recovery assessed at 15 days duration of arrest (p < 0.05) (Figure 4D).

**Figure 4.**
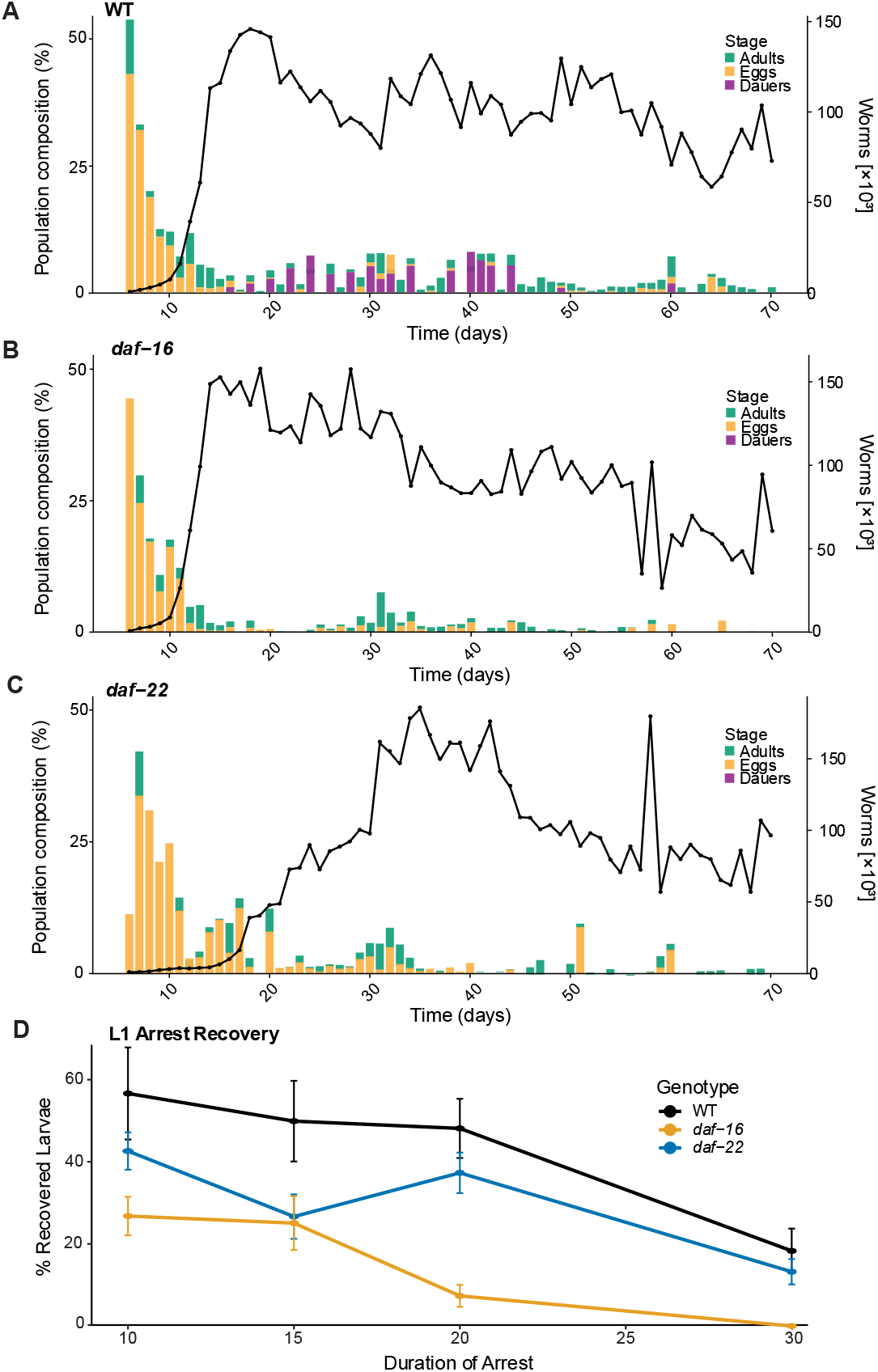
Stage distributions and diapause abilities within liquid culture populations differ between genotypes. (A, B, C) Time resolved stage distribution of populations with the population dynamics and associated total number of worms on the right axis. Bar graph colors denote developmental stages of individuals. Because larvae represented a majority of the population at most time points, the portion of the larval stage extending to 100% was left blank to allow clearer visualization of differences among other stages. Line graphs indicate average population dynamics calculated from independently maintained populations (A) WT (n=15 populations across three replicates) (B) *daf-22* (n=13 populations across three replicates) (C) *daf-16* (n=12 populations across three replicates). Time is shown in days on the x-axis (D) Recovery of individuals from L1 arrest based on duration of arrest in liquid media. Each datapoint represents the average recovery of a sample (n=3 replicates with 5-35 individuals scored per replicate). The recovery rate of arrested larvae is on the y-axis, and duration of L1 arrest is on the x-axis. Colors denote genotypes with error bars representing standard deviation.

## POPULATION DYNAMICS ARE AFFECTED BY NUTRIENT DEPRIVATION IN DIAPAUSE MUTANTS

To test whether genetic differences in developmental plasticity influence population recovery following an environmental stress, populations were exposed to a short period of food deprivation. Under short food deprivation conditions, where populations were culled and fed every 24 hours for 20 days followed by a deprivation period of no interference for 20 days and ending with a recovery phase of 20 days where populations were again culled and fed every 24 hours (Figure 5A), WT showed low disturbance in population dynamics (Figure 5B). *daf-22* had high disturbance to population dynamics with low population numbers after the deprivation phase but initiated population growth at reintroduction to food (Figure 5C). *daf-16* populations showed high disturbance to population dynamics with delayed and decreased population growth (Figure 5D). When compared to WT population dynamics, both *daf-22* and *daf-16* populations showed low similarity in the population dynamics distribution in PP-plots with *daf-22* (Figure 5E) exhibiting more similarity to WT populations than *daf-16* (Figure 5F). All WT populations survived the short deprivation, whereas extinction rates increased to 10% and 17% within *daf-22* and *daf-16* populations respectively (Figure 5G). WT populations exhibited a significantly higher average population size during the recovery phase (Figure 5H, Table 3) than either *daf-22* (p < 0.01) or *daf-16* (p < 0.0001). Additionally, *daf-22* had significantly smaller maximum population size in the recovery phase after a short deprivation when compared to WT (p < 0.05) (Figure 5I, Table 3). However, there was no reduction to maximum population size in *daf-16* population dynamics (Figure 5I, Table 3). These results indicate that WT populations have a distinct advantage over *daf-22* and *daf-16* populations when exposed to stress conditions.

**Figure 5.**
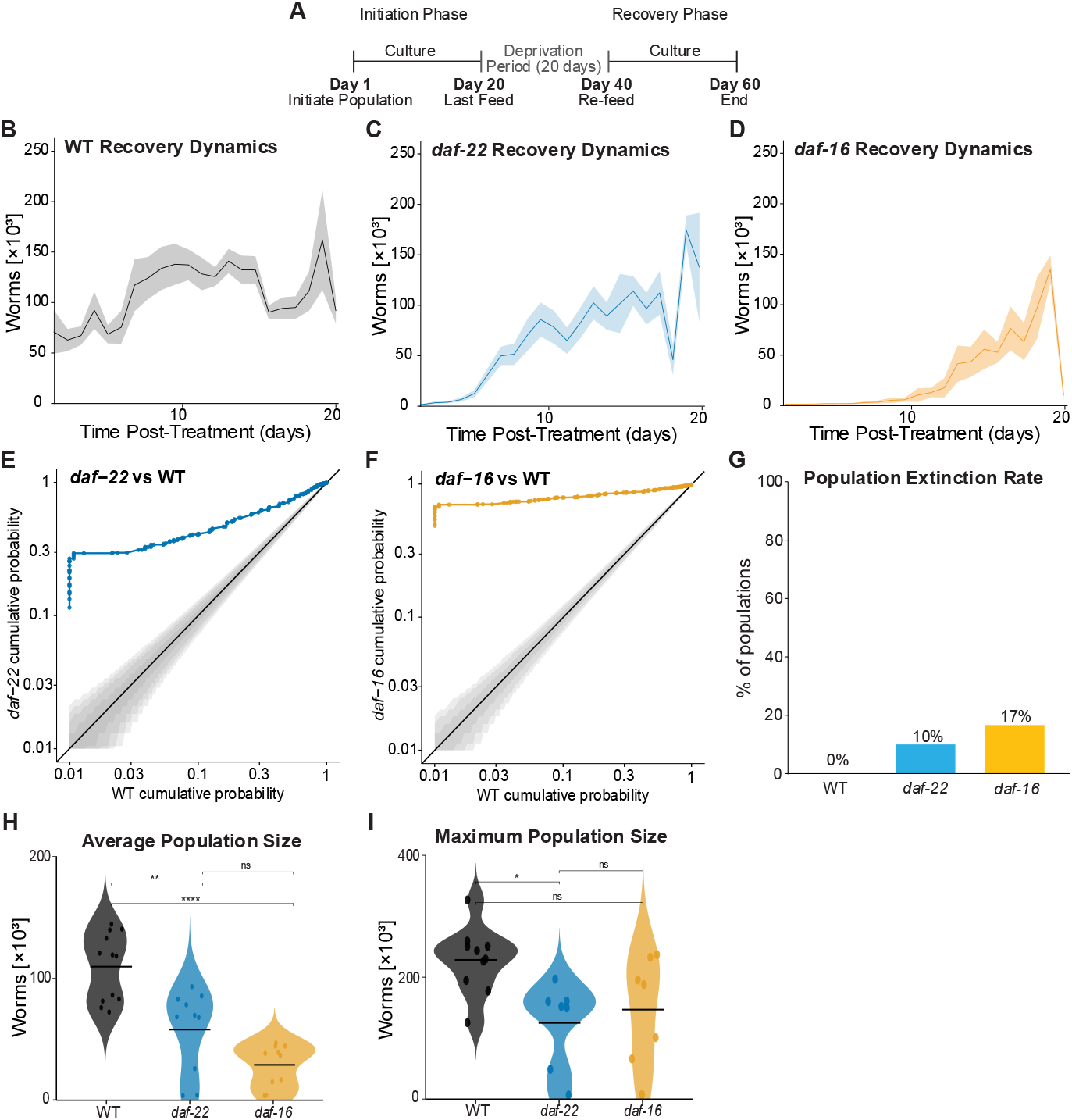
Population responses/ recovery dynamics to 20-day food deprivation phase. (A) Schematic of experiment conditions showing normal culture phase for 20 days, a deprivation phase of 20 days and a recovery phase of 20 days. All data shown is from the recovery period (days 40-60). (B, C, D) Population dynamics graph of WT (n=12), *daf-22* (n=9), and *daf-16* (n=10) mean total population size over time. Values represent mean ± S.E.M. across biological replicates, with technical replicates averaged prior to analysis. Population counts are expressed as worms x10^3^. (E, F) PP plots comparing empirical cumulative distribution functions of WT and mutant populations. Axes are in log scale, the black line indicates identical distributions, and the shaded region represents a 95% CI; deviations reflect differences in population dynamics. (G) Population extinction rates across genotypes under short stress conditions. Extinction was defined as 15 or more days with no detected worms in population counts post refeeding; the rate was defined as proportion of populations reaching extinction within each condition. (H) Average number of worms across genotypes from timepoints 40-60 shown as violin plots. Wilcoxon rank pairwise test for significance. (I) The maximum population size in the recovery period across genotypes shown as violin plots. The maximum was defined as the global maxima after the deprivation period. Wilcoxon rank pairwise test for significance. Significance: p< 0.05= *, p< 0.01=**, p< 0.001=***, p< 0.0001= ****; not significant = ns.

**Table 3.**
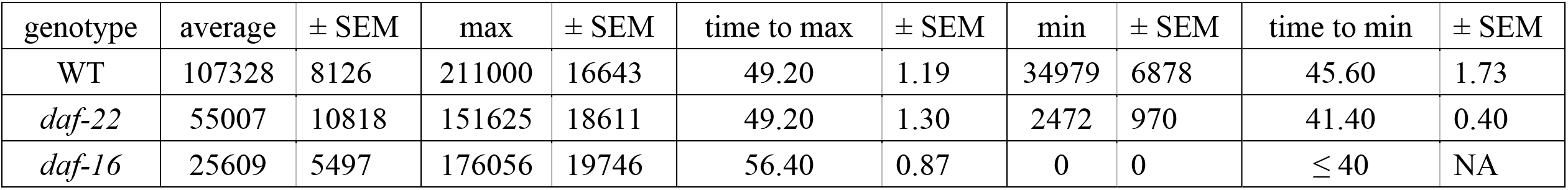
Population Summary Statistics under Short Nutrient Deprivation.

| genotype | average | ± SEM | max | ± SEM | time to max | ± SEM | min | ± SEM | time to min | ± SEM |
| --- | --- | --- | --- | --- | --- | --- | --- | --- | --- | --- |
| WT | 107328 | 8126 | 211000 | 16643 | 49.20 | 1.19 | 34979 | 6878 | 45.60 | 1.73 |
| *daf-22* | 55007 | 10818 | 151625 | 18611 | 49.20 | 1.30 | 2472 | 970 | 41.40 | 0.40 |
| *daf-16* | 25609 | 5497 | 176056 | 19746 | 56.40 | 0.87 | 0 | 0 | ≤ 40 | NA |

Under long food deprivation, where populations were culled and fed every 24 hours for 20 days followed by a deprivation period of no interference for 30 days and ending with a recovery phase of 20 days where populations were again culled and fed every 24 hours (Figure 6A), a similar pattern to short deprivation emerged, with stronger responses from the mutant genotypes. While WT populations showed minimal dynamics disturbance in response to the long deprivation phase (Figure 6B), both *daf-22* and *daf-16* populations started the recovery period with few to no live worms and failed to establish a population in the recovery period (Figure 6C, D). When comparing the distribution of population dynamics via PP-plot, *daf-22* populations showed low correlation to WT (Figure 5E) and *daf-16* populations showed no correlation to WT population dynamics in the recovery period (Figure 5F). All WT populations survived the 30-day deprivation period as well as the recovery period, where 50% of *daf-22* populations and 100% of *daf-16* populations went extinct (Figure 6G). As such, *daf-22* (p < 0.001) and *daf-16* (p < 0.1) had significantly reduced average population size during the recovery period when compared to WT populations (Figure 6H, Table 4).

**Figure 6.**
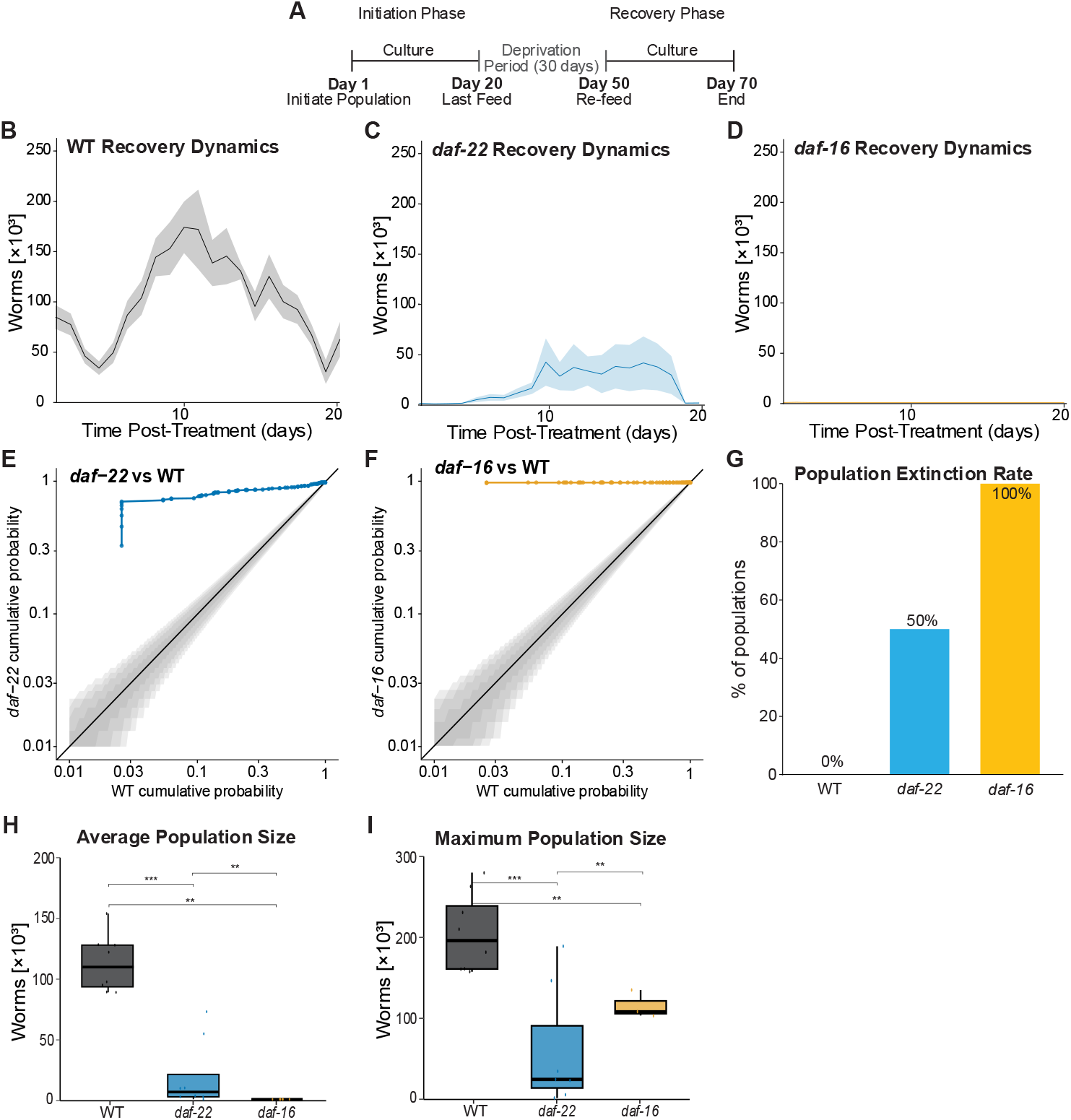
Population responses/ recovery dynamics to 30-day food deprivation phase. (A) Schematic of experiment conditions showing normal culture phase for 20 days, a deprivation phase of 30 days and a recovery phase of 20 days. All data shown is from the recovery period (days 40-60). (B, C, D) Population dynamics graph of WT (n=8), *daf-22* (n=8), and *daf-16* (n=7) mean total population size over time. Values represent mean ± S.E.M. across biological replicates, with technical replicates averaged prior to analysis. Population counts are expressed as worms x10^3^. (E, F) PP plots comparing empirical cumulative distribution functions of WT and mutant populations. Axes are in log scale, the black line indicates identical distributions, and the shaded region represents a 95% CI; deviations reflect differences in population dynamics. (G) Population extinction rates across genotypes under short stress conditions. Extinction was defined as 15 or more days with no detected worms in population counts post refeeding; the rate was defined as proportion of populations reaching extinction within each condition. (H) Average number of worms across genotypes from timepoints 40-60 shown as violin plots. Wilcoxon rank pairwise test for significance. (I) The maximum population size in the recovery period across genotypes shown as violin plots. The maximum was defined as the global maxima after the deprivation period. Wilcoxon rank pairwise test for significance. Significance: p< 0.05= *, p< 0.01=**, p< 0.001=***, p< 0.0001= ****; not significant = ns.

**Table 4.** Population Summary Statistics under Long Nutrient Deprivation.

| genotype | average | ± SEM | max | ± SEM | time to max | ± SEM | min | ± SEM | time to min | ± SEM |
| --- | --- | --- | --- | --- | --- | --- | --- | --- | --- | --- |
| WT | 112562 | 8393 | 205625 | 17054 | 59.60 | 1.24 | 30313 | 5003 | 53.4 | 1.02 |
| *daf-22* | 19049 | 9901 | 60643 | 28424 | 60.60 | 1.13 | 63 | 63 | 51.0 | NA |
| *daf-16* | 13 | 13 | 115500 | 9836 | NA | NA | 0 | 0 | ≤ 50 | NA |

Additionally, the maximum population size was significantly decreased in *daf-22* (p < 0.001) and *daf-16* (p < 0.01) populations during the recovery period (Figure 6I). These results indicate that mutations affecting the diapause ability of individuals impairs a population’s ability to survive and recover from harsh conditions.

## DISCUSSION

This study demonstrates that defects in starvation-responsive developmental arrest have limited effects on population dynamics under favorable conditions but become consequential under food deprivation. WT populations consistently showed greater resilience and persistence than either mutant strain, surviving both deprivation periods with minimal disruption. By contrast, *daf-22* and *daf-16* populations showed progressively reduced recovery and increased extinction as starvation duration increased. Although pleiotropic effects likely contributed to these phenotypes, the contrasting outcomes across genotypes support dauer formation and L1 arrest as key developmental strategies through which *C. elegans* populations persist during food limitation. Thus, genetic background modulates population-level responses most strongly when resources become limiting.

These outcomes broadly matched our expectations: WT was expected to persist under all conditions, *daf-16* to show the most severe phenotype, and *daf-22* to exhibit an intermediate response. WT animals retain the ability to enter L1 arrest when they hatch without food and dauer arrest under unfavorable conditions. *daf-22* mutants can enter L1 arrest and form dauers when supplied with exogenous pheromone, but they are defective in the biosynthesis of the density-dependent ascarosides required to coordinate dauer entry within a population (Golden & Riddle, 1985; Baugh & Sternberg, 2006; Butcher et al., 2009). Natural variation in ascaroside production across wild isolates, including a rare natural deletion encompassing *daf-22* that produces a pheromone-deficient phenotype similar to *daf-22*(ok693), further demonstrates that genetic variation in this pathway can alter chemical communication (Lee et al., 2023). In contrast, *daf-16* mutants are dauer defective and show impaired somatic developmental arrest and reduced survival during L1 starvation because they cannot execute the DAF-16/FOXO-dependent transcriptional response to reduced insulin-like signaling (Gottlieb & Ruvkun, 1994; Baugh & Sternberg, 2006; Kaplan et al., 2015). Stage distributions and arrest assays supported these expected phenotypes: SDS-resistant dauers were detected only in WT populations, whereas *daf-16* showed impaired recovery from extended L1 arrest (Figure 4). The altered dynamics of *daf-22* are therefore consistent with disrupted pheromone-mediated coordination of developmental decisions, whereas the more severe *daf-16* phenotype is consistent with broader impairment of developmental arrest and stress resistance.

Under control conditions, genotype affected the initiation and long-term trajectory of populations more than their established population size. Both mutant strains reached a larger first peak than WT, *daf-22* reached this peak later, and *daf-16* reached its post-peak maximum earlier; however, maximum, minimum and average population sizes were generally similar among genotypes once populations were established (Figure 2, 3, Table 1, 2). WT populations subsequently fluctuated around a stable mean, whereas *daf-22* and *daf-16* tended to decline after the first peak. This decline was accompanied by low background extinction in the mutants (9% for *daf-22* and 10% for *daf-16*), compared with no extinction in WT. Thus, diapause-related mutations altered aspects of population establishment and persistence even without imposed starvation, although most differences in population size were buffered under favorable conditions. Longer experiments would be needed to determine whether the declining mutant trajectories ultimately lead to greater extinction under otherwise favorable conditions.

Food deprivation exposed substantially stronger genotype-dependent differences. Following short deprivation, WT populations resumed their characteristic dynamics with little disturbance, whereas *daf-22* populations restarted growth from low abundance and *daf-16* populations showed delayed and reduced recovery (Figure 5). These differences became more pronounced after long deprivation: WT again recovered with little apparent impairment, while *daf-22* populations frequently failed to re-establish and *daf-16* populations did not recover (Figure 6). Extinction remained at 0% in WT across all conditions, but increased from 9% under control conditions to 10% and 50% after short and long deprivation, respectively, in *daf-22*, and from 10% to 17% and 100% in *daf-16*. The PP-plots similarly showed increasing divergence from WT with deprivation duration, with the greatest deviation in *daf-16* after long starvation (Figure 5, 6). Consistent with these trajectories, both mutants had lower average population sizes during recovery, while maximum population size declined in *daf-22* after both deprivation periods and in *daf-16* after long deprivation (Tables 3, 4). Together, these complementary measures show that starvation transformed relatively modest genotype differences into pronounced losses of population resilience and persistence.

These findings are consistent with previous studies showing that dauer-related pathways regulate not only individual developmental outcomes but also population growth, population size, dauer production and colony-level fitness (Green et al., 2013; Scharf et al., 2021; Chapman et al., 2024). More importantly, our results demonstrate that developmental arrest is not merely an individual response to starvation but a determinant of population persistence, resilience and extinction. WT populations survived both short-and long-term food deprivation with minimal disruption, consistent with the ability of *C. elegans* to exploit ephemeral microbial resources by alternating between rapid reproduction and starvation-resistant developmental arrest (Félix & Braendle, 2010; Baugh & Hu, 2020). Populations unable to enter either dauer or L1 arrest had little capacity to withstand food deprivation and invariably became extinct during prolonged starvation. Populations that retained L1 arrest but could not coordinate dauer formation recovered after short-term starvation, although more slowly and to lower abundance than WT, and half persisted even after prolonged starvation. Developmental plasticity and inter-individual communication therefore appear to stabilize populations across repeated cycles of growth and resource limitation.

The contrasting mutant outcomes also clarify the complementary contributions of L1 arrest and dauer. Larvae that hatch without food arrest at L1 and can remain viable for weeks, although survival and developmental recovery decline with arrest duration (Lee et al., 2001; Artyukhin et al., 2013; Baugh & Hu, 2020). L1-starved animals also communicate socially: higher larval density and soluble cues released during arrest can increase starvation survival, accelerate recovery and enhance DAF-16 activation (Artyukhin et al., 2013; Mata-Cabana et al., 2020). These findings provide a potential mechanistic context for the contribution of L1 arrest observed here and indicate that its protective capacity may depend not only on individual physiology but also on population density and chemical communication. Dauer larvae undergo more extensive morphological, metabolic and behavioral remodeling and can survive unfavorable conditions for months before resuming reproductive development when food returns (Golden & Riddle, 1984; Hu, 2007). Our results are consistent with dauer providing the strongest protection during prolonged starvation, but they also show that L1 arrest preserved half of the *daf-22* populations exposed to long food deprivation. L1 arrest may therefore be an underappreciated mechanism of long-term survival and temporal dispersal rather than only a transient response to early-life starvation. This finding challenges an exclusive emphasis on dauer as the dominant arrest stage under harsh conditions and instead suggests that WT persistence arises from complementary arrest programs operating over different developmental stages and environmental timescales.

Viewed more broadly, dauer and L1 arrest represent forms of temporal dispersal: rather than escaping unfavorable conditions spatially, individuals persist developmentally and re-enter the reproductive population when resources return. Comparable strategies occur across diverse taxa. Insects use diapause to synchronize active life stages with favorable periods (Denlinger, 2002, 2023; Sgrò et al., 2016); Daphnia and rotifers produce dormant eggs that form temporal reservoirs in sediments (Brendonck & De Meester, 2003; Franch-Gras et al., 2017); plant seed banks distribute germination across time and buffer populations against reproductive failure (Lennon et al., 2021; Yang et al., 2021); and dormant microbial cells and spores contribute to community recovery, persistence and diversity following nutrient limitation (Jones & Lennon, 2010; Bradley et al., 2025). Across these systems, dormancy delays reproduction but reduces the probability that all individuals encounter lethal conditions simultaneously. Our findings experimentally connect the loss of distinct arrest programs with whole-population outcomes, showing how disruption of individual developmental decisions can scale across generations to population extinction.

This connection may be increasingly important in variable environments, where the duration and predictability of resource limitation determine whether plastic responses protect populations or become mismatched to prevailing conditions (Reed et al., 2010; Sgrò et al., 2016; Bernhardt et al., 2020). The maintenance of both L1 arrest and dauer in WT *C. elegans* may broaden the range of conditions over which populations can persist: L1 arrest protects larvae encountering starvation immediately after hatching, whereas dauer enables later-stage larvae to withstand prolonged crowding and resource depletion. Multiple arrest strategies may consequently reduce fluctuations in the active population, preserve demographic and genetic reservoirs, and lower extinction risk across environmental timescales (Lennon et al., 2021; Webster et al., 2025). Here, loss of these strategies converted food deprivation from a temporary demographic disturbance into an elevated risk of population extinction.

Several limitations should be considered. These experiments used defined genetic backgrounds under controlled laboratory conditions, and outcomes may differ in genetically diverse natural populations or spatially and temporally heterogeneous environments. In addition, *daf-22* and *daf-16* mutations have pleiotropic effects on metabolism, reproduction and stress resistance, preventing the observed population phenotypes from being attributed exclusively to the absence of developmental arrest.

Future work should quantify the abundance and viability of arrested larvae throughout starvation, test genetic rescue or independent alleles, and compare laboratory WT with natural *C. elegans* isolates. Experiments incorporating fluctuating food availability, repeated starvation, additional environmental stressors and interspecific competition would further define when each arrest strategy promotes persistence. Together, these approaches will clarify how individual developmental decisions scale across generations to determine whether populations persist, recover or become extinct.

## CONCLUSIONS

Together these findings reveal that genetic regulation of developmental pathways and plasticity governs population level responses to environmental stress. The emergence of population differences of diapause-ability mutants under food deprivation underscores the importance of context in shaping biological outcomes. By linking genetic differences to population dynamics, this study advances our understanding of how conserved signaling pathways coordinate collective behavior in living systems. Both the volatility and the fluctuations of the diapause mutant populations suggest less regulation abilities compared to WT, likely stemming from the inability to enter diapause when appropriate and inter-individual communication. Pleiotropic effects of the mutations need to be investigated to elucidate effects seen as part of them may have been caused by interference to growth/development, metabolism, signaling etc. aside from diapause abilities. This suggests that genetic effects are not constitutive but are instead revealed under environmental stress. Such context-dependent phenotypic divergence is typical of stress-responsive regulatory pathways, including IIS and diapause. These findings are consistent with previous studies demonstrating that dauer-related pathways regulate not only individual developmental outcomes but also shape population-level dynamics. The observed differences between WT and dauer-ability mutants under food deprivation support that developmental plasticity and inter-individual communication play critical roles in population persistence, adaptation, and resilience.

## Acknowledgements

This work is part of GB’s master’s thesis “FROM INDIVIDUALS TO POPULATIONS: DRIVERS OF RESILIENCE” in the Department of Biological Sciences at Missouri S&T. We thank D. Duvernell and S. Minteer for their time and support as part of GB’s thesis committee. We thank K. Kornfeld for his support and scientific advice. We are grateful for the Caenorhabditis Genetics Center (funded by NIH Office of Research Infrastructure Programs, National Institutes of Health (P40 OD010440)) for providing strains. This work was supported in part by the NSF Award No. 2334884 under the project title “BRCBIO: The Impact of Pheromone Signaling on C. elegans Population Dynamics,”

## Author contributions

G.B. and A.S. conceived and designed the experiments. G.B., M.R., E.B., L.E., A.H., H.M. and A.S. performed experiments, G.B., H.M. and A.S. analyzed data, G.B, L.E. and A.S. provided scientific input, G.B. and A.S, wrote the article.

## Data availability

Data will be publicly available once the article is published.

## Code availability

Code will be publicly available once the article is published.

